# The tricellular junction protein ILDR2 in glomerulopathies: insights and implications

**DOI:** 10.1101/2024.05.03.592456

**Authors:** Florian Siegerist, Felix Kliewe, Elke Hammer, Paul Schakau, Joanne Ern Chi Soh, Claudia Weber, Victor Puelles, Maja Lindemeyer, Simone Reichelt-Wurm, Vedran Drenic, Christos Chatziantoniou, Christos E. Chadjichristos, Yiying Zhang, Miriam C. Banas, Matthias Nauck, Uwe Völker, Nicole Endlich

## Abstract

The tricellular tight junctions are crucial for the regulation of paracellular flux at tricellular junctions, where tricellulin (MARVELD2) and angulins (ILDR1, ILDR2 or LSR) are localized. The role of ILDR2 in podocytes, specialized epithelial cells in the kidney, is still unknown. We investigated the role of ILDR2 in glomeruli and its influence on blood filtration. Western blots, scRNA-seq and superresolution microscopy showed a strong expression of MARVELD2 and ILDR2 in podocytes that colocalizes with the podocyte-specific claudin CLDN5. Co-immunoprecipitation revealed that ILDR2 directly binds CLDN5. In glomerulopathies, induced by nephrotoxic serum and by DOCA-salt heminephrectomy, ILDR2 was strongly upregulated. Furthermore, *Ildr2* knockout mice exhibited glomerular hypertrophy and decreased podocyte density, however, did not develop effacement of podocyte foot processes or proteinuria. LC-MS/MS proteomic analysis of isolated glomeruli showed an increase in matrix proteins such as fibronectin and agrin. This suggests a protective role of ILDR2 in glomerulopathies.

## Introduction

Podocytes, a terminally differentiated cell type in the glomeruli, are essential for maintaining the integrity of the glomerular filtration barrier. These cells, characterized by interdigitating foot processes with filtration slits in between, are critical for the size selectivity of the filtration barrier, a property that is crucial for renal filtration function. In the past, the cell-cell contacts between podocytes have been described as specialized tight junctions^1^, with important regulatory proteins such as ZO-1 localized in these structures. Although podocyte junctions exist predominantly as bicellular connections, tricellular contacts have also been described, as shown by detailed reconstructions using scanning electron microscopy (SEM).^2^ At the site of these tricellular tight junctions (tTJs), specific proteins-such as tricellulin (MARVELD2) and the proteins of the angulin family (LSR, ILDR1 or ILDR2) are expressed.^3^ TTjs further influence the barrier function of epithelial and endothelial tissues, allowing the paracellular transport, e.g. of molecules and ions^4^ even independently of claudins.^5^ They could also play a mechanosensory role^6^ and will recruit claudins to the tTJs.^3^

Over decades, studies have shown that mutations in tTJ components are linked to a variety of diseases.^5^ Loss of tTJs components that govern tTJs homeostasis is known to disrupt the equilibrium of epithelium barrier function and monolayer organization, resulting in dysplasia. In 2021, Sugawara et al. also reported that aberrant expression angulin-1/LSR implicated the epithelial barrier function due to the loss of plasma membrane contact at the tTJs point. Furthermore, in 2017, a single cell RNA sequencing study identified ILDR2 as a podocyte-specific gene which was shown to be essential for the actin cytoskeleton in cultured podocytes.^7^ Recently, the group of O’Brian has shown that ILDR2 was associated with podocin along the slit diaphragm.^8^

This raised the question of where exactly ILDR2 is located in the glomeruli and what role ILDR2 plays in the podocytes. Using different methods and studying ILDR2 knockout mice, we investigated whether ILDR2 is essential for podocyte morphology and function as well as for the size selectivity of the filtration barrier.

## Material and Methods

### Generation of *Ildr2* KO Mice

The Ildr2-floxd line was generated as described by Millings et al.^9^ Whole body Ildr2 KO mice were generated by crossing Ildr2-floxed mice with B6.C-Tg(CMV-cre)1Cgn/J (JAX:006054). Mice were housed at room temperature in a 12-hr light/12hr-dark vivarium, with ad libitum access standard water. *Ildr2* knockout animals were generated by crossing animals with a floxed exon 1 of the *Ildr2* gene with a CMV:Cre line. The Cre was removed by backcrossing to the background strain (C57BL/6J). KO and WT animals were obtained by cross-breeding heterozygote animals mice. Experiments were performed with 6-24-month-old male and female mice. All studies were carried out in strict accordance with regulations in Germany regarding the use of laboratory animals.

### DOCA-salt induced hypertension

Induction of glomerular hypertension by uninephrectomy (UNX) and desoxycorticosterone acetate (DOCA) salt treatment has been described previously.^10^

### Mouse Model of Nephrotoxic Serum–Induced GN

Induction of NTS-induced glomerulonephritis has been described previously.^11,12^

### BTBR ob/ob mouse model of type 2 diabetes

Mice derived from the strain BTBR served as model for T2DM with BTBR WT mice as control and leptin-deficient BTBR ob/ob mice as diabetic animals. Heterozygous BTBR ob+/− mice (BTBR.Cg-Lepob/WiscJ, stock number: 004824; The Jackson Laboratory (Sacramento, CA, USA)) were used for breeding. All animal experimental procedures were approved by the local ethics committee at the University Hospital Regensburg (No.55.2-2532-2-1259).

### Immunofluorescence staining

For immunofluorescence of Ildr2 two polyclonal antibodies were used: 1.: An ILDR2 rabbit anti-mouse antiserum was a kind gift by Mikio Furuse (JCS 2015). Polyclonal antibodies were affinity-purified using the western-blot method with the immunization peptide. Fresh-frozen wildtype mouse kidneys were embedded in OCT (Sakura) and sectioned at 10 µm using a rotational cryomicrotome (HM500 Leica Microsystems, Wetzlar, Germany) and collected on super frost slides. Sections were air dried and fixed with 1:1 ethanol-acetone for 10 min at room temperature. Slides were washed in PBS, blocked, and primary antibodies incubated o/n at 4°C. Primary antibodies were detected using Alexa Fluor 488 conjugated goat anti-rabbit F(ab)2 fragments and Cy3 conjugated donkey anti-guinea pig full IgG antibodies. Nuclei were counterstained with DAPI and mounted in Mowiol for microscopy (Roth, Karlsruhe, Germany). 2.: Affinity purified rabbit anti ILDR2 antibody (ThermoFisher Scientific, Waltham, MA, U.S.A) diluted 1:150 in blocking solution was applied on 4 µm FFPE sections of mouse tissue after heat-mediated epitope retrieval in citrate buffer pH 9. For the detection of the primary antibody, the same protocol was used as for the cryosections.

### Single-mRNA in situ hybridization (RNAscope) and combined fluorescence *in situ* hybridization and immunofluorescence

*In situ* histochemistry was performed using the RNAscope Kits 2.5 Brown or Red with human and murine *ILDR2* exon 1-3, *PPIB* (positive control) and *dapB* (negative control) probes synthesized by ACDbio. Single chromogenic RNAscope was performed according to the manufacturer’s instructions with the following conditions: Heat mediated antigen retrieval for 15 min, and Proteinase Plus digestion for 30 min. hybridization of primary probe for 2 h at 40°C, AMP5 30 min at RT. The suitability of every FFPE block for RNAscope was evaluated using the PPIB positive control probe. Blocks with a *PPIB* score of 2 and higher went into further *ILDR2* testing.

For combined fluorescence *in situ* hybridization and immunofluorescence, we used the RNAscope Manual 2.5 Kit Red with AP and FastRed detection with the following modifications: After FastRed incubation, sections were washed in tap water and collected in 1x PBS. After blocking with 1% normal goat serum, 1% BSA, 1% FBS and 0.1% cold fish gelatin in 1x PBS, a pre-incubated mixture of affinity purified polyclonal rabbit anti podocin antibodies (IBL) 1:150 and Alex Fluor 488 conjugated site-specific dual monoclonal alpaca anti rabbit nano secondaries (single domain antibodies, VHHs, Chromotek) 1:1000 in blocking solution was applied to the sections at 4°C overnight. After 5 washes in 1x PBS and nuclear counterstain with DAPI, sections were mounted in Mowiol and imaged by confocal laser scanning microscopy using a Leica TCS-SP5 system using the 63x 1.4 NA oil immersion objective.

For semi-automated analysis of the ILDR2 expression, we established a FIJI macro that after manual selection of the podocin^+^ outlines of the glomerular tuft, segments, binarizes, and counts the *ILDR2*^*+*^ spots using the Particle Analyzer function in FIJI.^13^ Output values were: glomerular tuft area, ILDR2 particle number, mean and total area.

### Co-Immunoprecipitation and Western Blot

HEK-293 cells were cultured for 3 days and transfected with the respective myc or GFP plasmid using jetPEI (Polyplus-transfection, Illkirch, France). 48h post-transfection cells were scraped in ice cold PBS. Cells were washed twice in PBS and lysed in Pierce™ IP Lysis Buffer (Thermo Fisher Scientific) containing 1x Halt Protease and Phosphatase Inhibitor Cocktail (Thermo Fisher Scientific) for 20 min on ice. After removing debris by centrifugation for 20 min at 10,000 xg, cell lysates were incubated for 1.5h at room temperature with anti-myc-Trap Magnetic Beads (ChromoTek GmbH, Planegg-Martinsried, Germany). After two washing steps (10 mM Tris/Cl pH 7.5, 150 mM NaCl, 0.5 mM EDTA) beads were resuspended in 2 × SDS loading buffer (final concentrations: 64 mM Tris-HCl, 2% SDS, 10% glycerol, 0.1% bromophenol blue, 6.5 % 2-mercaptoethanol, pH 6.8) and boiled for 5 min at 95°C. Lysates were separated using a 4–20% Mini-PROTEAN^®^ TGX™ Gel, (Bio-Rad Laboratories, Munich, Germany) and transferred to a nitrocellulose membrane using a Trans-Blot^®^ Turbo^™^ Transfer System (Bio-Rad Laboratories) for 10 min at 2 A. Membranes were immersed for 1 h in blocking buffer (10 mM Tris, 100 mM NaCl, 5% non-fat dry milk, 0.2% Tween-20, pH 7.5) and incubated overnight at 4°C in TBS-Tween (0.5%) with the following antibodies: anti-ILDR2 (1:1000), anti-CLDN5 (1:1000), anti-CD9 (1:1000) and anti-tGFP (1:10000) for IP-WB. Blots were incubated for 1 h at RT with HRP–conjugated secondary antibody anti-mouse (SA00001-1, Proteintech Group) or anti-rabbit (SA00001-2, Proteintech Group) and developed using Clarity™ Western ECL Blotting Substrate (Bio-Rad Laboratories) and finally exposed to X-ray films (Fujifilm Super RX, FUJIFILM, Tokyo, Japan).

### RNA extraction, cDNA synthesis and qRT-PCR

Samples were processed in Tri-Reagent (Sigma-Aldrich) according to manufacturer’s instructions. For cDNA synthesis, 1 µg of the isolated total RNA was transcribed using the QuantiTect Reverse Transcription Kit (Qiagen, Hilden, Germany). The quantitative real-time PCR (qRT-PCR) analysis was performed on a QuantStudio™ 5 Real-Time PCR System (Thermo Fisher Scientific) using the iTaq Universal SYBR Green Supermix (Bio-Rad) with Gapdh as reference gene. Relative quantifications of the mRNA levels were done by the efficiency corrected calculation model by Pfaffl and are shown with standard deviations (SD) or standard error of the mean (SEM) from at least three biological replicates. Used primers can be viewed in Supplemental Table 1.

### Histology and Transmission Electron Microscopy

For routine histology, tissue was fixed in 4% paraformaldehyde pH 7.4 at RT o/n and embedded in paraffin after an ascending ethanol series. Sections were cut at 2-4 µm and collected on super frost slides (Marienfeld, Lauda-Königshofen, Germany).

For transmission electron microscopy, apical cortex blocks were trimmed to 1×1 mm and fixed in 4% glutaraldehyde in 1x PBS pH 7.4 for 2 days at RT. The tissue was embedded in EPON according to the manufacturer’s instructions. Semi– (500 nm) and ultrathin (70 nm) sections were cut using a Leica ultramicrotome and collected on glass slides and copper grids, respectively. Semithin sections were stained according to Richardson, and ultrathin sections were contrasted with lead-citrate.

### Liquid chromatography-mass spectrometry (LC-MS/MS)

Proteins were extracted from samples of five independent bioreplicates by heating glomeruli for 5 min at 95°C in 20 mM Hepes buffer containing 4 % SDS. Protein containing supernatant was collected by centrifugation (16.000 x g, 60 min, 4°C) and nucleic acid degraded enzymatically with universal nuclease (0.125 U/µg protein, (Pierce/Thermo, Rockford, IL, U.S.A.). Protein concentration was determined by BCA assay (Pierce/Thermo). Four µg protein were reduced by 2.5 mM dithiothreitol for 1 h at 60°C and alkylated with 10 mM iodoacetic acid for 30 min at 37°C in the dark before subsequent protein purification and proteolytic digestion according to a modified SP3 protocol.^14^ Peptides were separated by LC (Ultimate 3000, Thermo Electron, Bremen, Germany) before data-independent acquisition of MS data on an Exploris 480 mass spectrometer (Thermo Electron). MS data were analysed via the DirectDIA algorithm implemented in Spectronaut (v14, Biognosys, Zurich, Switzerland) using Uniprot database (v.2021-02) limited to Mus musculus (n=17063). Carbamidomethylation at cysteine was set as static modification, oxidation at methionine and protein N-terminal acetylation were defined as variable modifications, and up to two missed cleavages were allowed. Proteins were only considered for further analyses, if two or more unique+razor peptides were identified and quantified per protein. Further data analysis was performed as reported earlier.^15^ Detailed description of data acquisition and search parameters are provided in Supplemental table S2.

### Morphometry

Podocyte foot processes width in transmission electron micrographs were quantified by counting the number of filtration slits/glomerular capillary circumferences in FIJI.^13^ Processing, staining, imaging, and quantification were performed with minor modifications as described before.^16, 17^ In brief, 4 µm sections were directly mounted, blocked, and incubated with rabbit anti-podocin 1:150 (IBL International, Germany) and mouse anti-integrin α3 conjugated with Alexa Fluor 647 1:500 (Santa Cruz Biotechnology, Dallas, USA) and anti-rabbit Alexa Fluor 488-conjugated IgG 1:600 (ChromoTek, Germany) secondary antibody. Widefield and 3D-SIM images were acquired on an N-SIM super-resolution microscope (Nikon, Japan). To account for the 3-dimensional architecture of the filtration slit, 19 z-plane images were obtained to generate a maximum intensity projection (MIP) image. 3D-SIM images were reconstructed using NIS-Elements AR software (Nikon, Japan). Digital post-processing, profile plotting, and PEMP was performed using FIJI combined with a custom-built macro described before.^17^ First, in a whole slide image, glomeruli were segmented with an AI, and glomeruli with section artifacts or poor staining quality are excluded. The algorithm determined the total filtration slit length (FSL) based on podocin staining and the total podocyte foot process area based on integrin α3 staining (A). Filtration slit diaphragm density (FSD) was then calculated from the ratio of FSL over A.

### Microarrays on human kidney biopsies

Human renal biopsy specimens and Affymetrix microarray expression data were procured within the framework of the European Renal cDNA Bank – Kröner-Fresenius Biopsy Bank. Biopsies were obtained from patients after informed consent and with approval of the local ethics committees. Following renal biopsy, the tissue was transferred to RNase inhibitor and microdissected into glomeruli and tubulointerstitium. Total RNA was isolated from microdissected glomeruli, reverse transcribed, and linearly amplified according to a protocol previously reported.^18,19^

Previously generated microarray data from microdissected human glomeruli sourced from individuals with kidney disease and healthy donors were used (Affymetrix HGU133Plus2.0: GEO accession number: GSE47185 (H7-Glom: Affymetrix HGU133Plus2.0)). Pre-transplantation kidney biopsies from living donors were used as control (GEO accession number: GSE37463 (H7-Glom: Affymetrix HGU133Plus2.0)). CEL file normalization was performed with the Robust Multichip Average method using RMAExpress (version 1.20) and the human Entrez-Gene custom CDF annotation from Brain Array (version 25). To identify differentially expressed genes, the SAM (Significance Analysis of Microarrays) method was applied using SAM function in Multiple Experiment Viewer (TiGR MeV, Version 4.9).^20^ A q-value below 5% was considered to be statistically significant. Analysis included gene expression profiles from patients with diabetic nephropathy (DN; n=7), minimal change disease (MCD; n=5), focal segmental glomerulosclerosis (FSGS; n=10) and rapidly progressive glomerulonephritis (RPGN, n=23) as well as controls (living donors, n=18).

### Confocal laser scanning microscopy

Images were captured with an Olympus FV3000 confocal microscope (Olympus, Tokyo, Japan) with 20x/40x/60x oil immersion objectives and Olympus FV3000 CellSense software.

### Statistical analysis

The GraphPad Prism 9 software was used for statistical analysis of experimental data and preparation of graphs. Scatter plots indicate individual units used for statistical testing (samples, cells or replicates), as specified in the respective figure legends. Data are given as means ±SD or ±SEM, analyzed by unpaired t test with repeated measurements (n).

For multiple groups statistical analyses were done by ANOVA followed by a Benjamini-Hochberg post-hoc test. Statistical significance was defined as p < 0.05 and significance levels are indicated as * p < 0.05, ** p < 0.01, *** p < 0.001, **** p < 0.0001 or non-significant (ns). The number of independent experiments and analyzed units are stated in the figure legends.

## Results

### Podocytes express the tricellular tight junction-associated protein ILDR2

Tricellular junctions are a universal feature of epithelial cells and contain tricellulin (MARVELD2) which is bound to at least one angulin (ILDR1, ILDR2 or LSR). In the Sampson Normal Tissue microarray dataset of the Nephroseq database (http://www.nephroseq.org) derived from microdissected human kidneys, ILDR2 was enriched in the glomerular fraction in comparison to other kidney fractions (Fig 1A). Single-cell RNA sequencing of murine kidneys showed that in comparison to the other angulins, Ildr2 was enriched within the podocyte cluster together with Marveld2 and the podocyte-specific claudin, claudin-5 (Fig. 1B). We verified these findings by RT-PCR of murine glomeruli isolated using the dynabead perfusion method, which showed glomerulus-enriched expression of *Ildr2* as well as a weaker expression of *Marveld2* (Fig. 1C). This finding was quantified with RT-qPCR which showed that *Ildr2* was around 15-fold enriched in isolated glomeruli in comparison with total kidney fractions (Fig. 1D).

**Figure 1:**
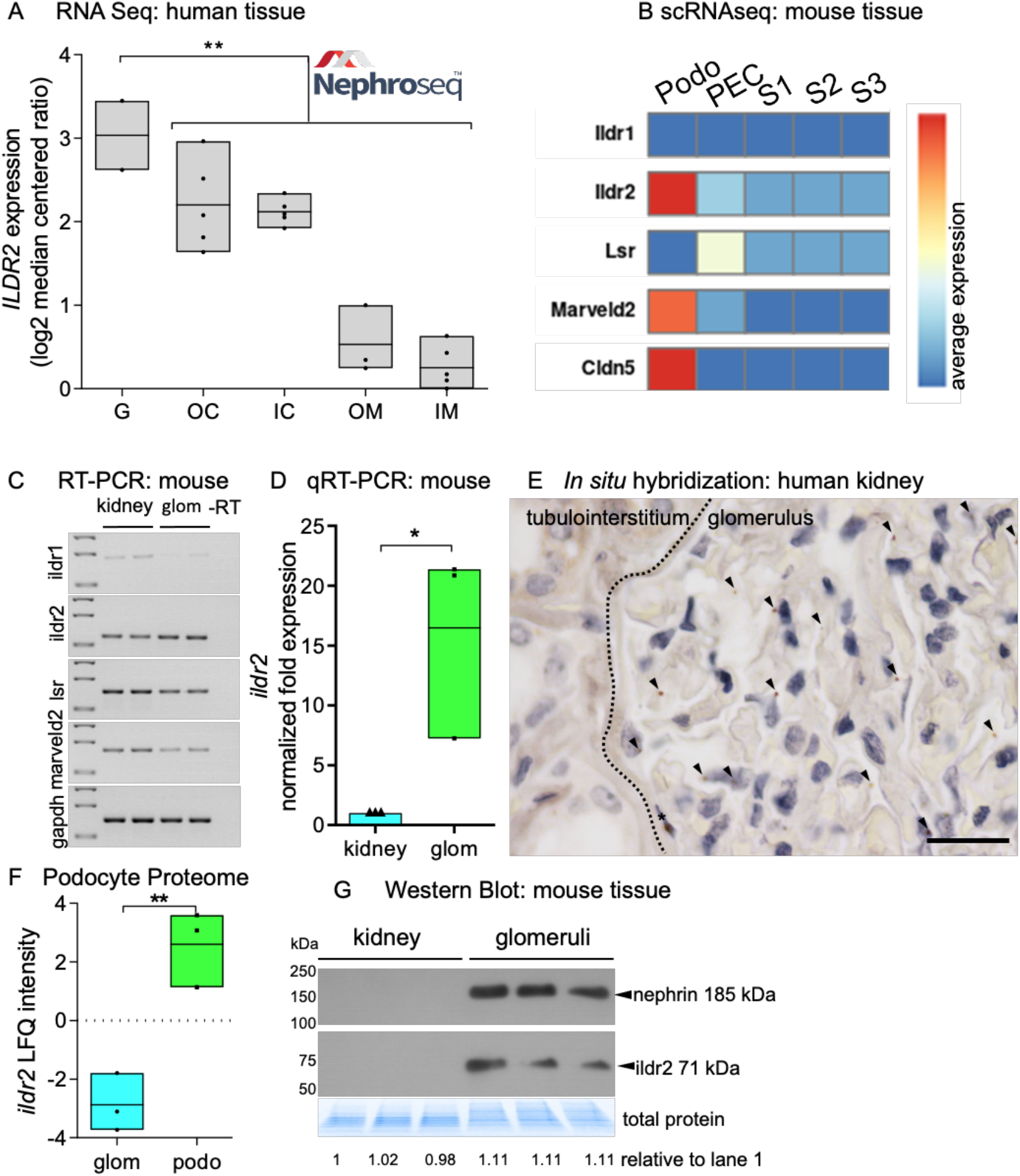
*ILDR2* expression in human and mouse kidney tissue. (A) *ILDR2* expression in glomeruli (G), outer renal cortex (OC), inner renal cortex (IC), outer renal medulla (OM) and inner renal medulla (IM). Data are represented as log2 median centered ratio. Each dot represents one sample. Data were taken from www.nephroseq.org. (B) Database analysis of *Ildr2* in podos (podocytes), PECs (parietal epithelial cells) and PT (segment 1, 2 or 3 of proximal tubule) using the KidneyCellExplorer (Ransick et al., 2019) based on a single-cell RNA sequencing data set of murine kidneys.^24^ (C) RT-PCR of *Ildr1, Ildr2, Lsr* and *Marveld2* in murine kidney and isolated glomeruli. Gapdh served as a loading control. (D) The mRNA expression of *Ildr2* in isolated glomeruli and murine kidneys was quantified by qRT-PCR (n=3). qRT-PCR experiments were normalized to the kidney samples and Gapdh. (E) *In situ* hybridization of *ILDR2* in human kidney tissue. Scale bar represents 20 µm. (F) Podocyte proteome data showed an enrichment of ILDR2 protein level in podocytes compared to glomeruli. Data from Rinschen et al..^25^ (G) Western Blot confirmed glomerulus-specific expression of ILDR2 (n=3). Nephrin served as positive control and total protein served as a loading control. * p<0.05; ** p<0.01.

Chromogenic single-RNA *in situ* hybridization using the RNAscope method showed the specific presence of *Ildr2* mRNA exclusively in podocytes in healthy human and mouse kidney tissue sections (Fig. 1E). In a published isolated podocyte versus bulk glomerulus mass spectrometry dataset, podocyte-specificity of ILDR2 in comparison to whole glomeruli could be verified (Fig. 1F). Anti-ILDR2 western blots of isolated glomeruli (purity indicated by the presence of nephrin) versus the whole kidney fraction were performed. Glomerular lysates probed with a polyclonal anti-ILDR2 antibody showed a single band of around the predicted size of 71 kDa (Fig. 1 G).

Immunofluorescence staining using a specific antibody against ILDR2 detected an epitope in murine and human glomeruli which produced a punctate staining pattern (Fig. 2A, Fig. S1). Confocal laser scanning micrographs (C-LSM) showed that within glomeruli, ILDR2-spots co-localized with nephrin, a marker of the podocyte slit diaphragm in mouse and human tissue sections (Fig. 2A).

**Figure 2:**
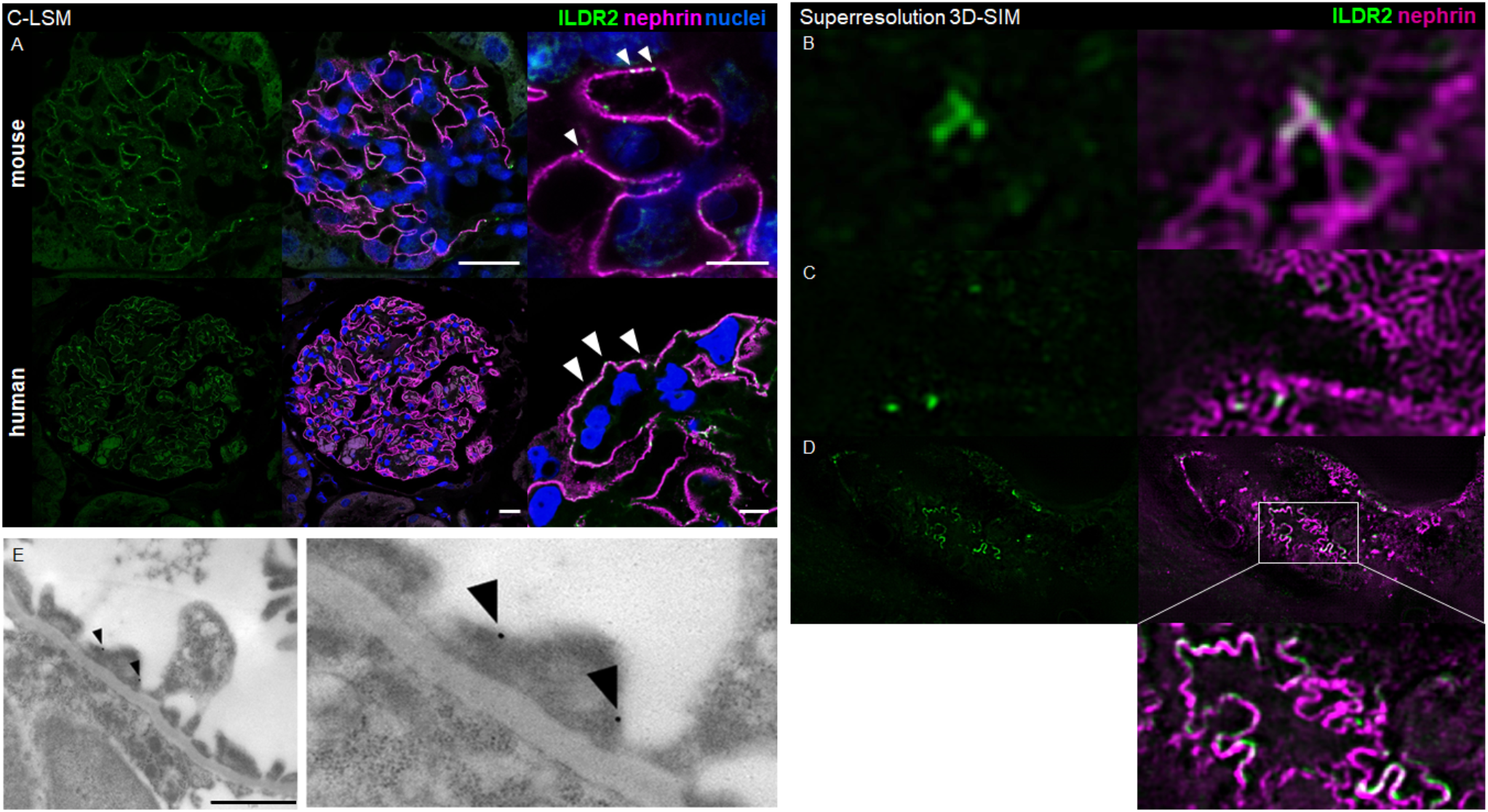
Podocytes express the tricellular tight junction-associated protein ILDR2. (A) Immunofluorescence staining for ILDR2 (green) and the podocyte-specific marker nephrin (magenta) in mouse and human kidney tissue. Nuclei were stained with Hoechst (blue). Scale bars represent 20 µm and 5 µm (magnification), respectively. (B, C) Super-resolution 3D-structured illumination microscopy (3D-SIM) revealed that ILDR2 (shown in green) localized to tricellular junctions as indicated by a co-staining with nephrin (shown in magenta) on Y-intersections of the nephrin-stained filtration slit and to bicellular junctions in a spotted manner within healthy-appearing foot processes. (D) Immunofluorescence staining of kidney sections of E19 embryonic mouse kidneys showed that ILDR2 broadly localized to effaced podocyte filtration slits as indicated by a linearized double ILDR2 (green) and nephrin (magenta) cell-cell contact. (E) Immunogold electron microscopy showed cell-membrane-associated binding of the anti-ILDR2 antibody podocyte foot processes. Scale bar represents 1 µm.

Super-resolution 3D-structured illumination microscopy (3D-SIM) revealed that ILDR2 localized at tricellular junctions as indicated by co-staining with nephrin on intersections of the nephrin-stained filtration slit (Fig. 2B). Additional to this, focal ILDR2 localized to some bicellular junctions in a punctated manner. These ILDR2-positive junctions were associated within normally formed foot processes (Fig. 2C). Immunogold transmission electron microscopy (TEM) validated this observation. Here we found ILDR2 localized to the cell membrane of podocyte foot processes (Fig. 2E).

Looking in embryonic mouse kidneys at E19 by 3D-SIM, ILDR2 was broadly localized to the developing filtration slits (Fig. 2D). The areas where nephrin and ILDR2 colocalized were more frequent and broader compared to healthy adult sections. In areas where podocyte foot processes were broadened (effaced), ILDR2 colocalized to the filtration slits in a linear and colocalized manner as shown by 3D-SIM (Fig. 2D).

### ILDR2 directly binds the podocyte-specific Claudin-5

CLDN5 has been shown to be the podocyte-specific claudin in bicellular tight junctions which is recruited and increased during the development of podocyte foot process effacement^9^. Using co-immunoprecipitation (co-IP) of HEK293T cells that were co-transfected with ILDR2-myc and CLDN5-tGFP, we found a direct interaction of both proteins (Fig. 3A). In double-transfected cells, Cldn5-tGFP could be detected in anti-myc pulldown eluates, whereas single-transfected cells did not show a signal. Additionally, analysis of immunofluorescence stainings by SR-SIM showed that the expression of ILDR2 was co-localized with CLDN5 in human kidney glomeruli in areas of podocyte foot process effacement (Fig. 3B).

**Figure 3:**
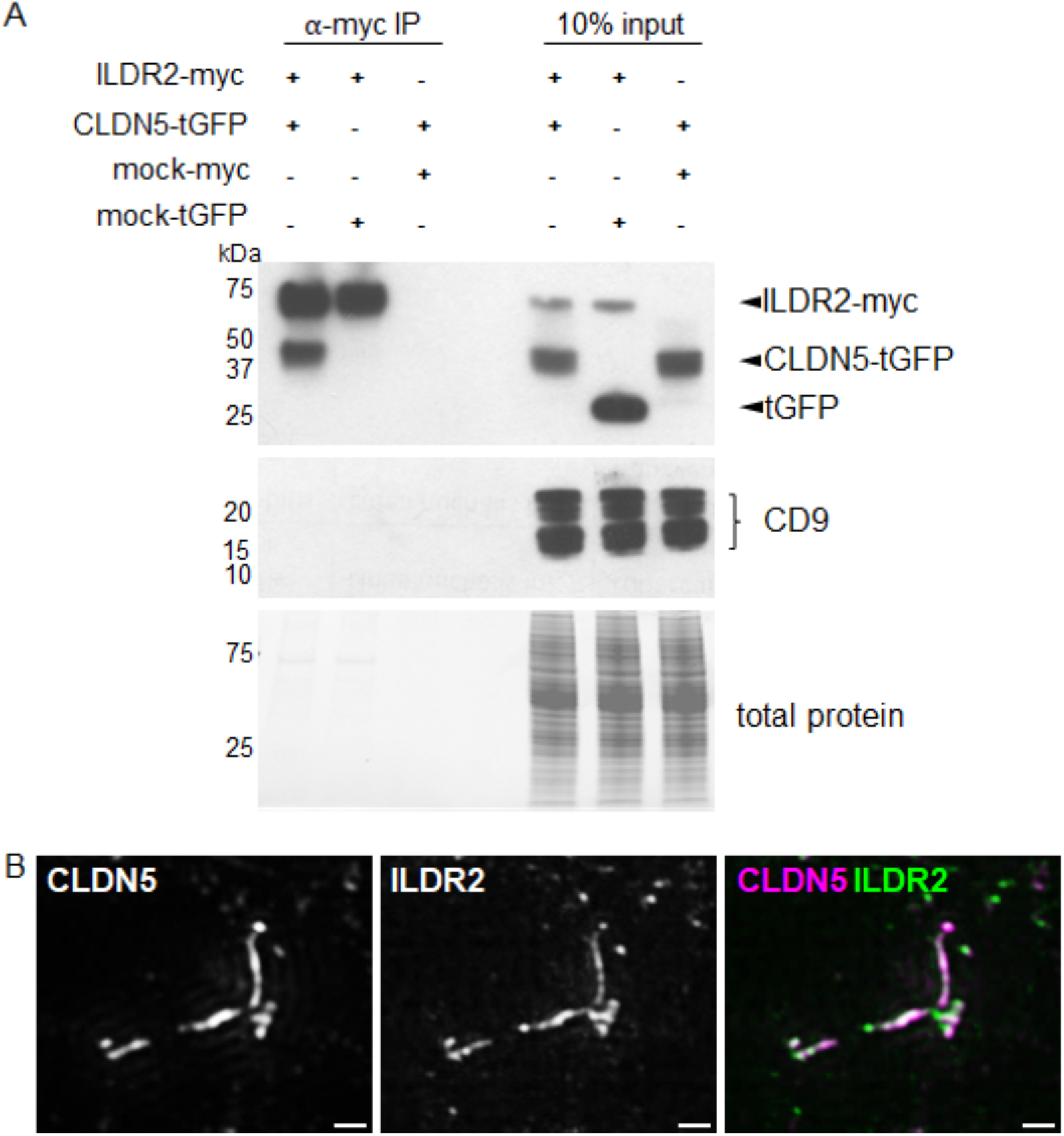
ILDR2 is an integral component of podocyte bi– and tricellular tight junctions bound by the podocyte-specific claudin 5. (A) Co-immunoprecipitation experiments of ILDR2-myc and CLDN5-GFP followed by Western blot analysis with indicated antibodies. “Input” means the sample on 10% of volume used for IP. Anti-CD9 Western blot served as a (non-binding) negative control. (B) The expression of ILDR2 (green) co-localized with CLDN5 (magenta) in human kidney glomeruli in areas of podocyte foot process effacement. Imaged by SR-SIM. Scale bars represent 500 nm.

### ILDR2 is up-regulated in different glomerulopathies in mice and human

To find out the role of ILDR2 *in vivo*, we analyzed different murine models of glomerular disease. In mice which were treated with nephrotoxic serum to induce crescentic glomerulonephritis as well as in the DOCA salt/UNX mice, a model to mimic hypertension-induced glomerular damage, we found an upregulation of the protein level as well as mRNA by immunofluorescent staining and *in situ* hybridization. (Fig. 4A-F).

**Figure 4:**
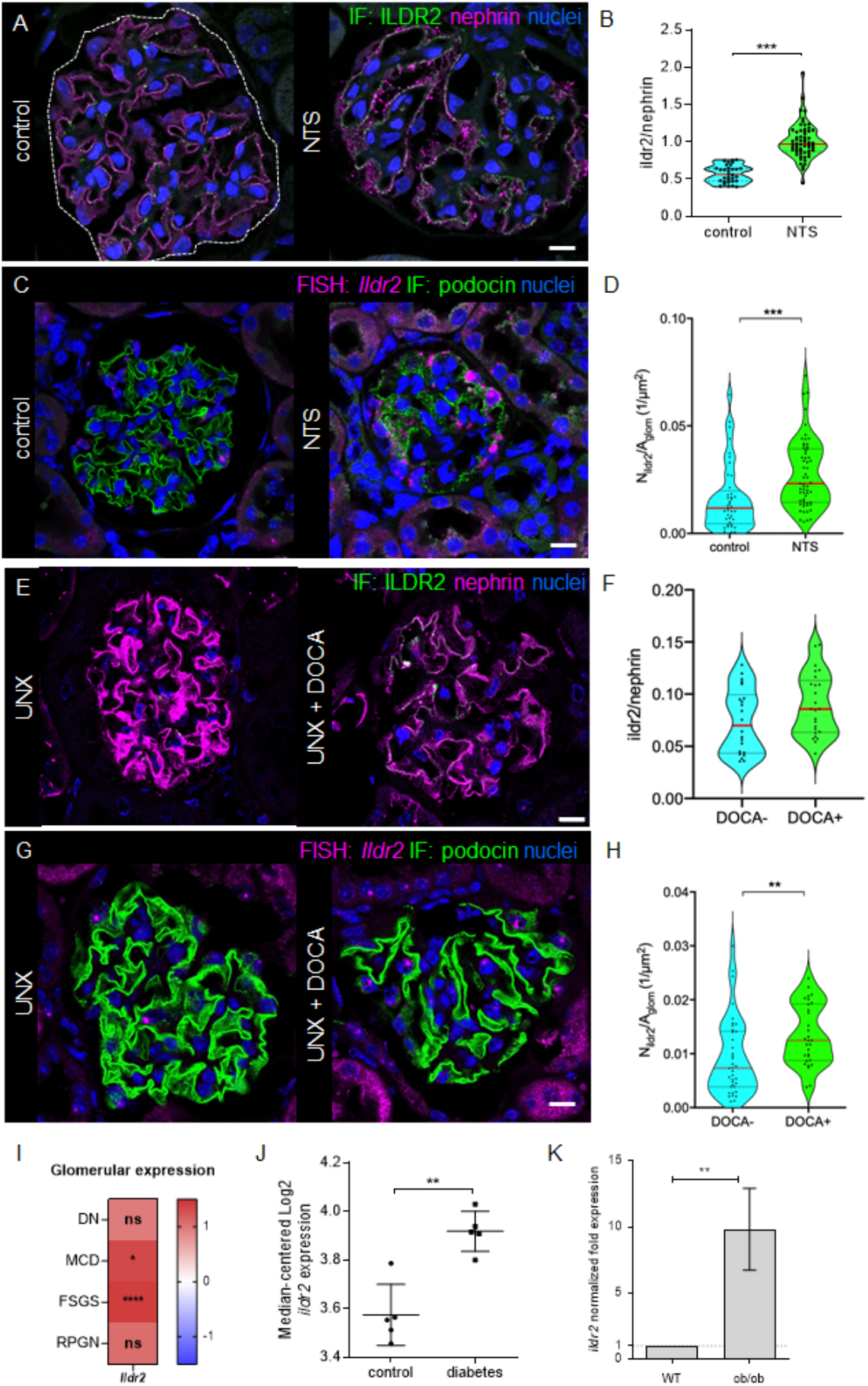
ILDR2 is up-regulated in DOCA-salt/ UNX and NTS-treated mice as well as in different human glomerulopathies. ILDR2 expression (shown in green) in control and NTS-treated mice (A) or uninephrectomy (UNX) and desoxycorticosterone acetate (DOCA)-salt-treated mice (E). Nephrin (magenta) served as a podocyte-specific marker. NTS injected mice (B) and DOCA-salt treated (F) mice showed significantly increased ildr2/nephrin ratio. RNAScope of *Ildr2* (magenta) followed by quantification of ILDR2 positive spots per glomerulus area (1/µm^2^) in NTS-treated mice (C, D) and DOCA-salt-treated mice (G, H) confirmed the increased expression of *Ildr2* after injury. Podocin (shown in green) served as a podocyte marker. (I) mRNA expression level of microdissected glomeruli from renal biopsies of human patients suffering from diabetic nephropathy (DN; n=7), minimal change disease (MCD; n=5), focal segmental glomerulosclerosis (FSGS; n=10) and rapidly progressive glomerulonephritis (RPGN, n=23) compared with healthy living donors (LD; n=18). Data are represented as log2 fold change, a q-value <0.05 was considered as significant. Red: up-regulated to LD. (J) Analysis of diabetes mouse glomeruli (Nephroseq database) substantiated increased expression on mRNA levels of *Ildr2* in diabetes mice (n=5) compared to Ctrl (n=5). (K) An increase in the *Ildr2* expression was also observed in ob/ob mice (n=18). Data are presented as means± SD (J), means ± SEM (K) or violin plot (B, D, F, H); * p<0.05; ** p<0.01; *** p<0.001; **** p<0.0001 ns, not significant. Scale bars represent 10 µm.

Furthermore, *Ildr2* mRNA levels were significantly upregulated in diabetes mouse glomeruli (Fig. 4J) and in 24 weeks old diabetic ob/ob mice compared to wildtype control (Fig. 4K). To compare the expression of *ILDR2* between glomerulopathies and healthy individuals, we analyzed microarray data from glomeruli from kidney biopsies of patients diagnosed with DN, MCD, FSGS and RPGN and found a significant increase of *ILDR2* expression in the glomeruli of MCD and FSGS patients (Fig. 4I).

### *Ildr2* knockout mice do not develop proteinuria or changes of podocyte foot processes

*Ildr2* knockout animals (*Ildr2* KO) were generated by crossing animals with a floxed exon 1 of the *Ildr2* gene with a CMV:Cre line (Fig. 5A). The Cre was removed by backcrossing to the background strain (C57BL/6J). Offspring were produced in the expected mendelian ratios and developed without obvious phenotypic changes. The efficient knockout of *Ildr2* was verified with RT-PCR and RT-qPCR which showed no significant amplicon (Fig. 5B and C). Both Western blot and immunofluorescence staining showed no binding of the anti-ILDR2 antibody in the kidneys of knockout animals (Fig. 5D and Fig. S2A). While the general histologic analysis of the kidneys of these animals was without obvious alterations, in-depth morphometric analysis with transmission electron microscopy (TEM) revealed partial podocyte foot effacement (Fig. 5F). Analysis by podocyte exact morphology measurement procedure (PEMP) showed a slightly, but not significantly reduced filtration slit diaphragm density in *Ildr2* KO mice compared to WT mice (3.98 µm^-1^ versus 4.31 µm^-1^) (Fig. 5H). Model-based glomerular morphometry showed slight glomerular hypertrophy with reduced podocyte density (Fig. 5I-K). However, besides these alterations in total glomerular volume (Fig. 5K) as well as podocyte foot process morphology (Fig. 5F), animals did not show significantly elevated albuminuria in comparison to respective wild-type littermates (ACR in µg/mg: WT: 15; KO: 42) (Fig. 5L).

**Figure 5:**
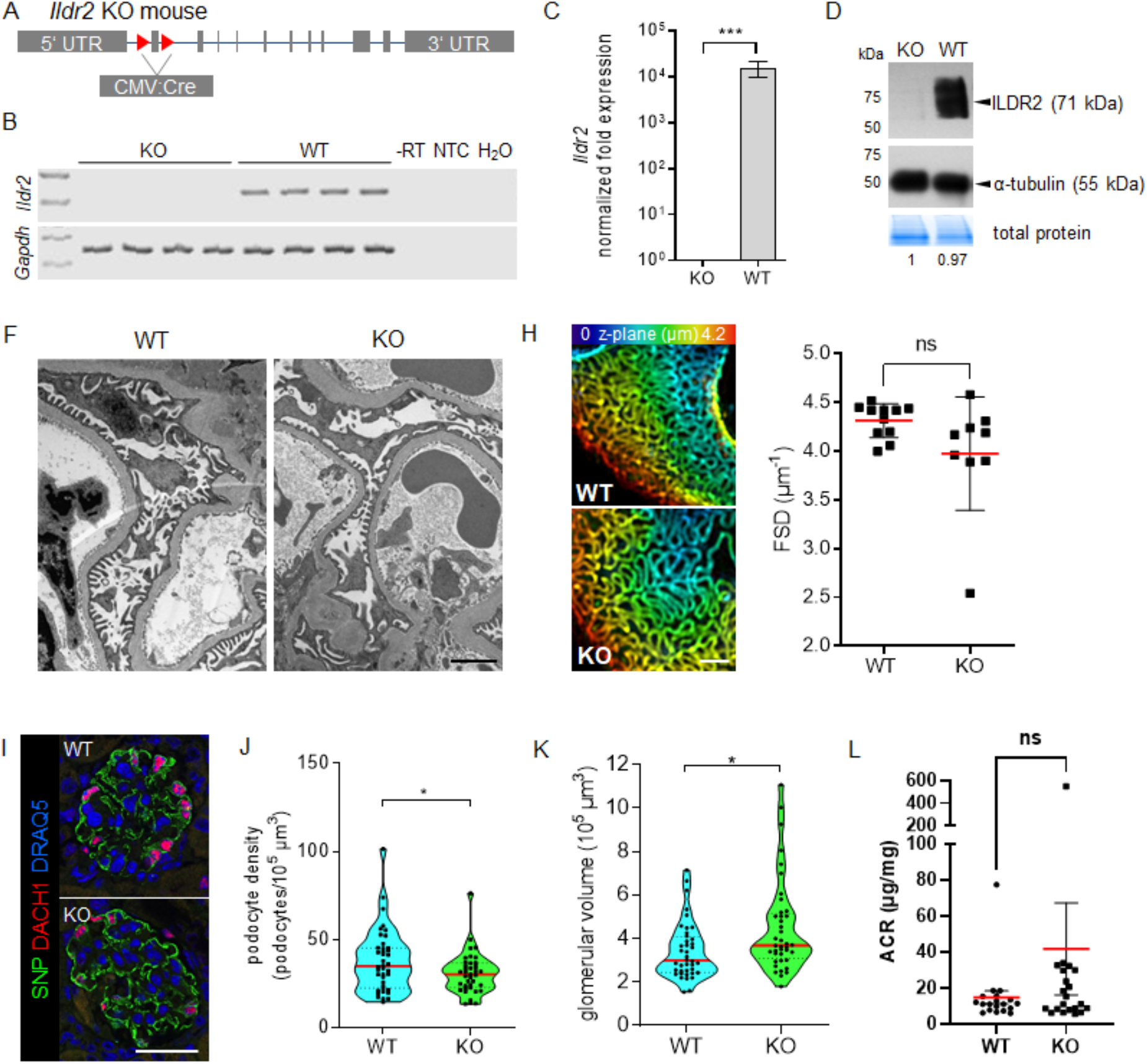
*Ildr2* knockout mice do not develop proteinuria but have altered podocyte morphology. (A) Schemata of *Ildr2* whole-body knockout animals by crossing animals with a floxed exon 1 of the ILDR2 gene with a CMV:Cre line. (B) *Ildr2* KO verification by RT-PCR. (C) *Ildr2* KO verification by qRT-PCR. Gapdh served as a reference. (D) *Ildr2* KO verification by Western Blot.Alpha-Tubulin and total protein served as a loading control. (F) Transmission electron microscopy of WT and *Ildr2* KO mice. (H) 3D-SIM in *Ildr2* KO animals showed no significantly reduced filtration slit density (FSD) indicative for aberrant FP architecture. SIM data of 11 WT and 9 KO animals were quantified by PEMP (at 24 months of age). Kidney sections were stained for the slit diaphragm protein podocin. Z axis scales of 3D-SIM were color coded as indicated. (I) mRNA expression level of microdissected glomeruli from renal biopsies of human patients suffering from diabetic nephropathy (DN; n=7), minimal change disease (MCD; n=5), focal segmental glomerulosclerosis (FSGS; n=10) and rapidly progressive glomerulonephritis (RPGN, n=23) compared with healthy living donors (LD; n=18). Data are represented as log2 fold change, a q-value <0.05 was considered as significant. Red: up-regulated to LD. (L) Urinary albumin-creatinine ratio measurements indicated no significantly increased levels of proteinuria in *Ildr2* KO mice in comparison to WT animals (each individual dot represents one experimental animal). Scale bars represent 2 µm (F), 1 µm(H), 50 µm (1).* p<0.05; ** p<0.01; **** p<0.0001; ns, not significant.

### *Ildr2* KO mice show glomerular profibrotic alterations, but no compensatory up-regulation of angulin or other tight junction proteins

To find out which other proteins are regulated by the loss of ILDR2, we performed proteomic profiling of isolated glomeruli from five independent WT and five *Ildr2* KO mice by LC-MS/MS. We detected 128 significantly up-and 46 down-regulated proteins in the *Ildr2* KO mice compared to WT (p < 0.05 and fold change > 1.25) (Fig. 6A). Proteomic data impressively showed that extracellular matrix (ECM) proteins such as agrin and fibronectin were significantly up-regulated in *Ildr2* KO glomeruli (Fig. 6B).

**Figure 6:**
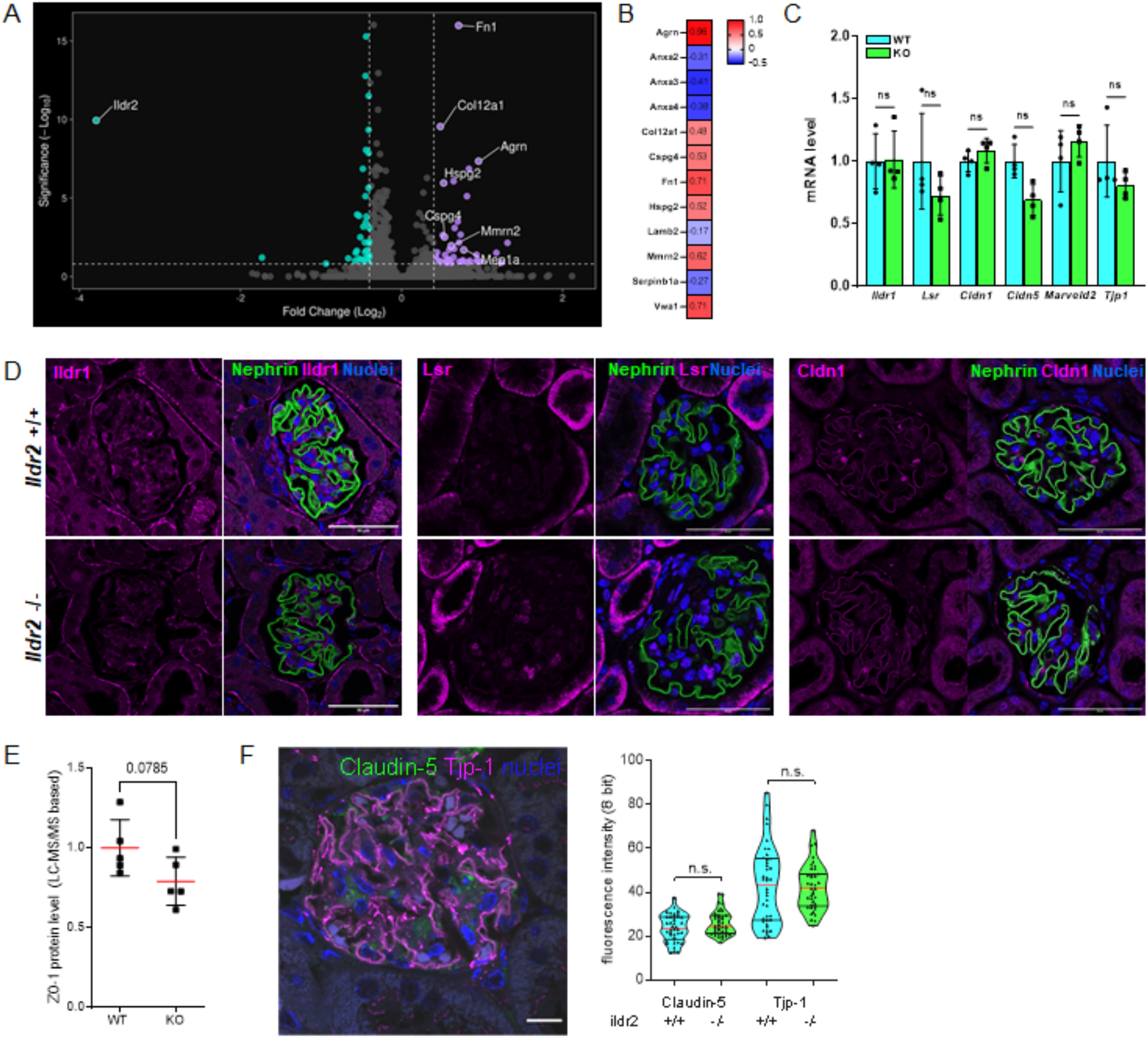
*Ildr2* KO glomeruli showed increased expression of extracellular matrix proteins, but no up-regulation of potentially compensatory angulin or tight junction proteins. (A) Volcano plot displaying proteins with significantly changed abundance in *Ildr2* KO mice in comparison to wildtype mice (n=5). Cyan: down-regulated compared to controls; purple: up-regulated compared to controls. Given are proteins with a fold-change ≥ 1.25 and p-value ≤ 0.05. (B) Ratios of regulated extracellular matrix proteins in *Ildr2* KO mice. Data are represented as log2 fold change, a p-value <0.05 was considered as significant. Red: up-regulated; blue: down-regulated compared to controls. (C) The mRNA expression of *lldrl, Lsr, Cldnl, Cldn5, Marveld2* was not significantly regulated in *Ildr2* –I-mice (n=4). qRT-PCR experiments were normalized to wildtype mice (WT), *Gapdh* served as a reference, ns, not significant. (D) Immunofluorescence staining for lldrl, Lsr and Cldn1 (red) and the podocyte-specific marker nephrin (shown in green) in *Ildr2* KO (Ildr2 –/-) and WT mice (*Ildr2* +/+). Scale bars represent 50 µm. (E) LC-MS/MS based relative protein level of ZO-1 (TJP1) in isolated glomeruli from *Ildr2* KO and *Ildr2* WT mice (n=5). (F) Fluorescence intensity measurement of immunofluorescence stained glomeruli with specific antibodies against claudin-5 and Tjp1 (ZO-1). Scale bar represents 10 pm.

Interestingly, at the protein level, there was no increase of other angulins such as ILDR1 or LSR in the *Ildr2* KO animals. Using qRT-PCR, we analyzed the mRNA levels of *Ildr1, Cldn1, Cldn5, Lsr* and *Marveld2*. None of them showed significant changes due to the loss of *Ildr2* (Fig. 6C). This was also confirmed by immunofluorescence stainings (Fig. 6D-F & Fig. S2B-C). The tight junction protein ZO-1 also showed no significant regulation at mRNA or protein level (Fig. 6C, E). Furthermore, no increased albumin uptake was detected in the renal tubules (Fig. S2D).

## Discussion

Podocytes are important for proper blood filtration and form a specialized cell-cell junction between their interdigitating foot processes. Since the seminal paper of the Farquhar group, it has been known that this cell-cell contact contains tight junction features.^1^

As it has been demonstrated that podocytes form tricellular junctions already under healthy conditions,^2^ we wanted to investigate which proteins of the tricellular tight junction complex were expressed at these junctions. Here we show that the angulin ILDR2 is expressed at tricellular tight junctions under healthy conditions which is in agreement with a recent publication shown by Higashi et al..^21^ We further observed, that after the onset of glomerulopathies, its localization shifted from tricellular to bicellular tight junctions. We observed a similar pattern in the expression of podocyte-specific claudin 5, whose distribution shifted from a bicellular punctate to a linear pattern.^12^ This raised the question about how ILDR2 was integrated within the slit diaphragm. Using co-immunoprecipitation, we could show a direct interaction of ILDR2 and claudin 5, suggesting that ILDR2 is integrated within the slit diaphragm as part of the tight junction complex. Therefore, the tri-to bicellular translocation directly follows the elongation of the filtration slit and disease-associated dedifferentiation of the podocyte cell-cell contact. Therefore, direct interaction with podocin is rather unlikely and in the aforementioned work by Gerlach and colleagues, ILDR2 has been pulled-down by podocin as a part of the large multiprotein slit diaphragm complex.^8^

To reveal whether ILDR2 was essential for the integrity of the size selectivity of the filtration barrier, we phenotyped ILDR2 whole-body knockout mice. After validation of the loss of the ILDR2 expression in podocytes, we analyzed the urinary creatinine/albumin ratio. Surprisingly, we have not found any significant changes of the foot processes. However, we have observed an increase of the glomerular size, indicating hyperfiltration with consecutive glomerular hypertrophy.

The precise function of tricellular tTJs in regulating paracellular permeability across epithelia has been a subject of investigation. In 2017, Krug demonstrated in an *in vitro* assay that tTJs facilitate up to 29% of paracellular macromolecule flux, in particularly tight epithelial cell monolayers.^22^ Conversely, in cells with more permeable tight junctions, the contribution of tTJs to flux is rather minimal.

Consequently, it can be postulated that in podocytes, which experience significant paracellular flux due to glomerular filtration, and therefore are a rather leaky epithelium, the regulatory role of tTJs in permeability may be less significant compared to tight epithelia. This is in line with Higashi et al., who in 2013 demonstrated that ILDR2 provided a rather weak transepithelial barrier in comparison to the other angulins when transfected in epithelial cell monolayers.^3^

Therefore, while it is recognized that tTJs can influence the passage of smaller macromolecules across epithelia, their impact on macromolecular selectivity within the glomerular filtration barrier appears to be rather limited.^23^ Suggesting that depletion of ILDR2 resulted in a gross disruption of ILDR2-containing tTJs, this could be an explanation why *Ildr2* KO animals were not proteinuric, as the baseline flux of macromolecules like albumin across tTJs appears to be negligible under steady-state conditions.

Proteomic profiling of isolated glomeruli of *Ildr2* KO animals revealed that the *Ildr2* KO had a significant impact on different signaling pathways. We have identified the up-regulation of several extracellular matrix proteins indicating a pro-fibrotic glomerular phenotype.

Since no change of the size selectivity of the filtration barrier occurred in *Ildr2* KO animals, we wanted to find out whether one of the other known angulin proteins was locally up-regulated to rescue the ILDR2 loss in podocytes. However, we have not found an up-regulation of ILDR1 or LSR. Besides these candidates, we have not detected other tight-junction-associated proteins that could be responsible for a rescue of the loss of ILDR2.

In summary, our results show that ILDR2 is an important component of the glomerular filtration barrier, with increased expression and localization observed in various glomerulopathies. While loss of ILDR2 does not change podocyte foot process morphology, it affects podocyte number and matrix expression, suggesting to be a counterplayer of profibrotic events during glomerulopathies.

## Acknowledgements

This work was supported by a start-up grant of the Research Network Molecular Medicine and by a grant of the Gerhard-Domagk Masterclass of the University Medicine Greifswald to Florian Siegerist and by grants of the Federal Ministry of Education and Research (BMBF, grant 01GM1518B; STOP‐FSGS and grant 01ZX1908B; 01ZX2208B, Sys_CARE) to Nicole Endlich and Uwe Völker. Maja Lindenmeyer is supported by BMBF, grant 01GM2202A; STOP‐FSGS and BMBF grant 01EK2105D, UPTAKE. We thank all participating centers of the European Renal cDNA Bank-Kroener-Fresenius Biopsy Bank (ERCB-KFB) and their patients for their cooperation. Active members at the time of the study are listed in ref. (Martini et al JASN 2014, Vol 25, No 11, 2559-2572). This work was generously supported by the Südmeyer Stiftung für Nieren– und Gefäßforschung and the Dr. Gerhard Büchtemann fund, Hamburg, Germany. The funders had no role in study design, data collection and analysis, decision to publish, or preparation of the manuscript.

## Author Contributions Statement

The study was designed by FS and NE. FS and CW contributed to the cell culture experiments; biopsies were handled and analyzed by FS, PS, JS and ML; LC-MS/MS was performed by EH and UV; animal experiments were performed by FS, CW, SRW, MB, CC and CEC; Western blots and IP were performed and quantified by FS, CW and FK.; VD performed PEMP; all other experiments were performed by FS; experimental data were analyzed by FS, FK, PS and VP. MN analyzed urine. FS, FK and NE wrote the main manuscript text. FS and FK prepared figures. All authors reviewed the manuscript.

## Data availability

All data supporting the findings of this study are available within the paper and its Supplementary Information. Further information is available from the authors upon reasonable request.

## Ethics declarations

The authors declare no competing interests.

## Figures & Figure Legends

**Figure S1:**
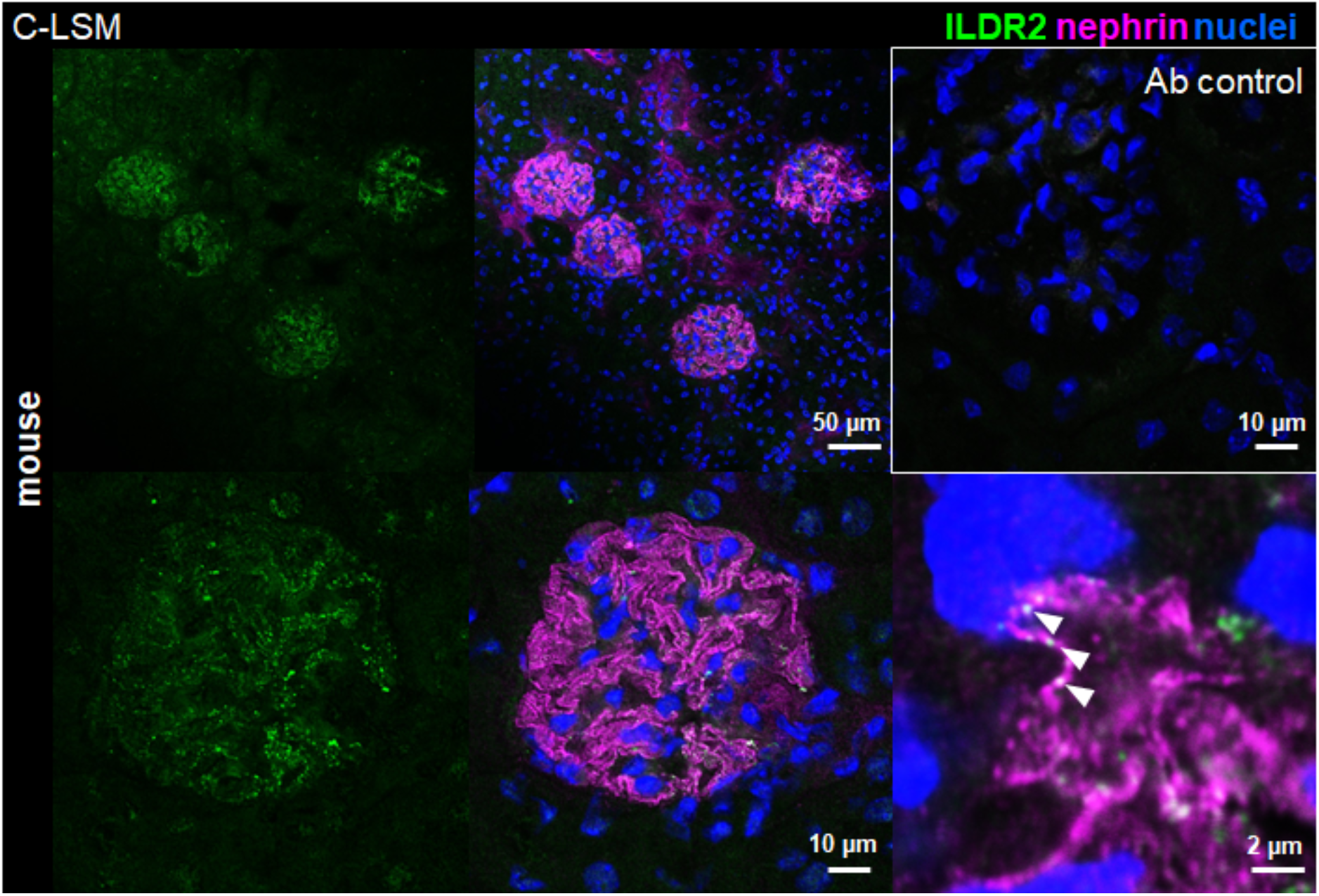
Expression of the tricellular tight junction-associated protein ILDR2. Immunofluorescence staining for ILDR2 (green) with the antibody from Mikio Furuse and the podocyte-specific marker nephrin (magenta) in mouse kidney tissue. Nuclei were stained with Hoechst (blue).

**Figure S2:**
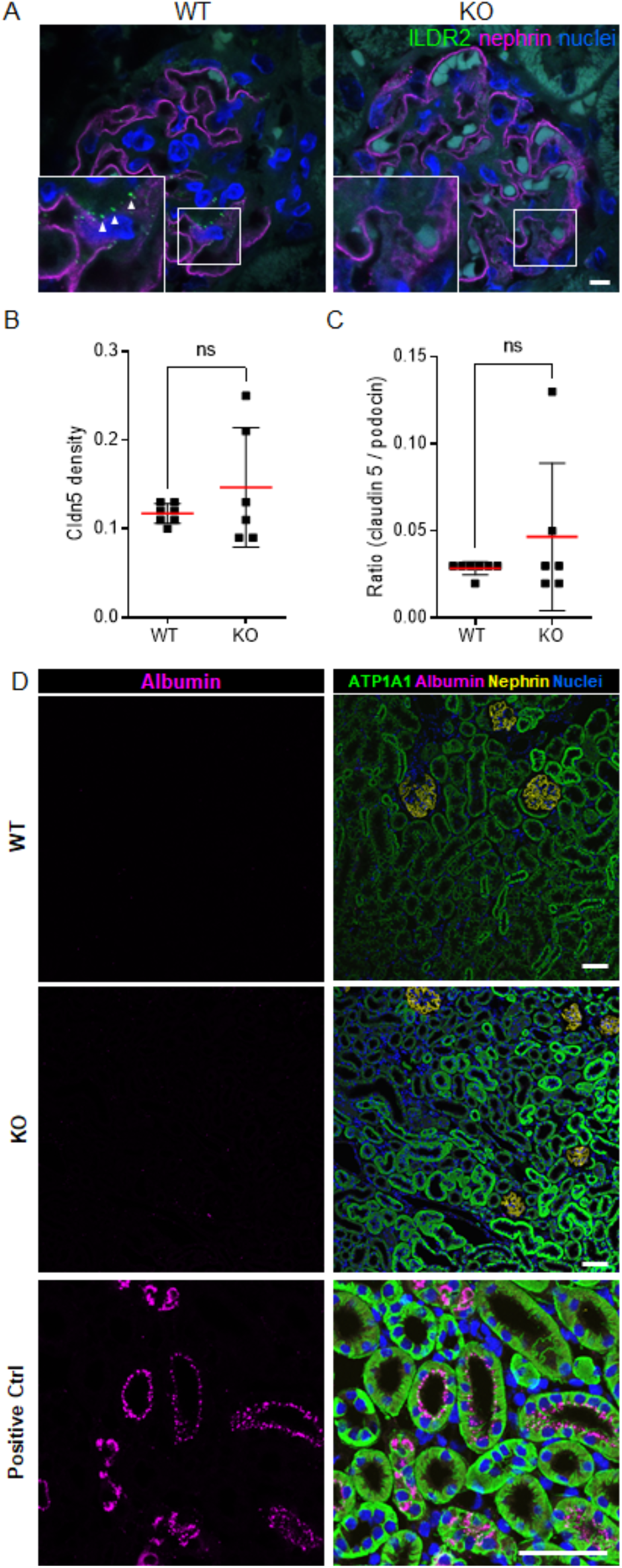
*Ildr2* KO mice showed no regulation of Cldn5 and no increased albumin uptake in proximal tubules. (A) Immunofluorescence staining for ILDR2 (green) and the podocyte-specific marker nephrin (magenta) were used to validate efficient deletion of *Ildr2* in *Ildr2* KO mice in contrast to *Ildr2* wildtype (WT) mice. (B) Claudin-5 (Cldn5) density in WT and Ildr2 KO mice. (C) Claudin-5 / podocin ratio in WT and Ildr2 KO mice. (D) Immunofluorescence staining for albumin (magenta), ATP1A1 (green) and the podocyte-specific marker nephrin (yellow) in *Ildr2* KO and WT mice. NTS-treated mice served as positive control. Nuclei were stained with Hoechst (blue). Scale bars represent 5 µm (A) and 50 µm (D).

